# Functional characterization and classification of mechanosensitive bladder afferents

**DOI:** 10.64898/2026.05.27.728257

**Authors:** Guadalupe Manrique-Maldonado, Xuejiao Sun, Stephanie L. Daugherty, Jonathan M. Beckel, Marcelo D. Carattino

## Abstract

Normal urinary bladder function relies on afferent fibers that detect and integrate mechanical and chemical cues related to bladder distension. Though, the molecular identity and function of the various sensory neuron types involved in bladder function have yet to be fully elucidated. Here, we introduce a novel framework for the functional classification of mechanosensitive bladder afferents based on their differential responses to physiological (15 μl/min) and noxious filling (30 s at intravesical pressures of 10, 20, 30, 40, 50, and 60 cmH₂O). Our data reveal the presence of three distinct types of mechanosensitive bladder afferents, two that respond to physiological distension (type I and II) and one that is activated by noxious stimulation (type III). Of the two populations that respond to physiological filling, one displays a linear increase in firing with bladder filling (type I), while the firing of the other plateaus as intravesical pressure increases (type II). Fast filling (130 μl/min) increases the discharge of all three afferent types, with the effect being most pronounced in those responding to noxious stimulation (type III). Corroborating the existence of three functionally distinct bladder afferent populations, Yoda1, a selective PIEZO1 channel activator, significantly increased the firing rate of types I and III during slow filling and of type III during noxious stimulation. In summary, we present a reliable and reproducible method for studying and classifying bladder afferents, while providing compelling evidence for the existence of functionally distinct populations of mechanosensitive afferents, each activated and regulated by distinct mechanisms.

**New & Noteworthy:** Using a novel approach, we identify three types of mechanosensitive afferents innervating the urinary bladder, two that respond to slow filling and one that is activated only by noxious distension. The three afferent types display distinct firing patterns during rapid filling and in response to the PIEZO1 channel agonist Yoda1.

## Introduction

Urine storage and micturition are controlled by a central neural circuit that receives input from afferents nerve fibers embedded in the mucosal and muscular layers of the bladder. These bladder-innervating sensory neurons are pseudo-unipolar, with one axonal branch terminating in the bladder wall and another entering the spinal cord and forming synapses with second-order neurons in the dorsal horn and dorsal gray commissure (1–3). Sensory neurons are responsible for detecting chemical and mechanical stimuli in the bladder wall and transmitting this information to higher neural centers. However, afferents not only transduce signals related to bladder distension but are also integral to pathological states, where they function as nociceptors encoding noxious stimuli. Despite decades of study, the molecular identities and functional roles of the various sensory neuron subtypes innervating the urinary bladder as well as the mechanisms by which they sense physiological and pathological processes and communicate information to central neurons have not been fully elucidated. Establishing a relationship between the physiological responses and molecular identities of distinct afferent populations will be essential for targeting them therapeutically.

Bladder afferents have been studied using nerve recordings in *ex vivo* and *in vivo* preparations (4–22). These studies have revealed the existence of several bladder afferent populations with distinct functional properties, consistent with both the pronounced heterogeneity reported using immunohistochemical methods (23–28) and the large transcriptional diversity observed in single-cell and single-nuclei RNA-seq analyses of dorsal root ganglia (DRG) (29–32). Afferents that respond to bladder filling *in vivo* or *ex vivo* recordings have been classified based on the threshold pressure for activation (e.g., low or high threshold) and conduction velocity (i.e., Aδ fibers or C fibers) (4–12, 16, 21, 22). For bladder sheet preparations, afferents have been classified according to their responses to mechanical stimuli applied to the receptive field, including stretch and mucosal and serosal stroking with von Frey filaments (14, 15, 17–20). While these studies have advanced our understanding of bladder afferent function and regulation, the physiological roles and mechanisms underlying mucosal and serosal afferent activation remain unclear, and translating their responses to physiological *in vivo* conditions is not straightforward.

The consensus is that micturition depends on mechanosensitive, thinly myelinated, Aδ afferent fibers that respond to bladder distention in the physiological range (23, 33). In contrast, unmyelinated C fibers have high mechanical thresholds and respond to bladder distention only at elevated pressures (4–6). Because capsaicin, a TRPV1 agonist, thought to be expressed only in C fibers, does not affect normal micturition reflex in cats and rats, it has been inferred that C fibers are not required for voiding (1, 34–36). However, a population of C fibers that respond to bladder distention in the physiological range of pressures has also been reported (7–11, 37). And indeed, recent transcriptomic analyses of DRG neurons indicate that TRPV1, a non-selective cation channel activated by capsaicin, is expressed in both peptidergic (CGRP-positive) neurons and in non-peptidergic neurons that express neurofilament heavy (NFH), a marker of myelinated fibers (30). Thus, the traditional view that Aδ afferent fibers mediate normal bladder function is likely oversimplified, underscoring the need to reevaluate the functional diversity of bladder afferent populations.

Here, we use multi-unit pelvic nerve recordings and an *ex vivo* preparation to study the mechanical properties and regulation of murine bladder-innervating afferents. We developed a systematic approach to analyze and classify bladder afferents based on their response to both physiological and noxious filling. Using this framework, we identified three populations of bladder afferents: two activated at physiological filling pressures and one that responds to noxious mechanical stimulation. These three populations of bladder-innervating afferents exhibit distinct responses to physiological, fast and noxious distension as well as to the PIEZO1 channel activator Yoda1. The implications of our findings for bladder physiology are discussed.

## Materials and methods

### Mice

All experimental procedures were approved by the University of Pittsburgh Institutional Animal Care and Use Committee. C57BL/6J mice were obtained from the Jackson Laboratory (Bar Harbor, ME). Mice were housed in standard cages at the University of Pittsburgh under 12h light/12h dark cycles with free access to food and water. Experimental mice were 2 to 5 month-old virgin females and group housed up to 5 per cage. The stage of the estrous cycle was not monitored.

### Ex vivo bladder preparation

Mice were euthanized by CO_2_ inhalation followed by a thoracotomy and transcardiac perfusion with cold Krebs solution containing (in mM): NaCl 118.5, KCl 4.7, NaHCO_3_ 25, NaH_2_PO4 1.3, MgSO4 1.2, D-Glucose 18, and CaCl2 2.5. All tissues rostral to the kidneys were removed and the preparation was transferred to a 15-cm dish lined with Sylgard and filled with cold Krebs solution. A cannula was inserted in the urethra, and the ureters were tied close to the urinary bladder wall. Kidneys, ovaries, and uterine horns were carefully removed. A silk ligature was placed around the sciatic nerves at the level of the leg bone to expose the pelvic nerves around the L6-S1 vertebrae and L6 and S1 spinal roots were cut close to their exit from the intervertebral foramen. At this point, the bladder, urethra and associated spinal nerve roots were removed and carefully transferred to a recording chamber equilibrated with warm Krebs solution bubbled with a mixture of 95% O_2_/5% CO_2_ (v/v). The solution in the chamber was continuously recirculated, and the temperature maintained at 35°C with an inline heater (Warner instruments, model SH-27B) coupled to a dual automate temperature controller (Warner instruments, model TC-344B). The urethral cannula was connected to a four-way connector: one branch led to a pressure transducer (World precision instruments, Cat. No. BLPR2), a second port was connected to a syringe pump (World Precision instruments, model SP100iZ) for continuous infusion and to a pressure stand used of noxious filling with Krebs, and the third port to a cannula with a pressure release valve, which facilitated emptying the bladder after filling (Fig.1A). A 3-way stopcock valve was used to switch between syringe pump and pressure stand filling.

**Figure 1.**
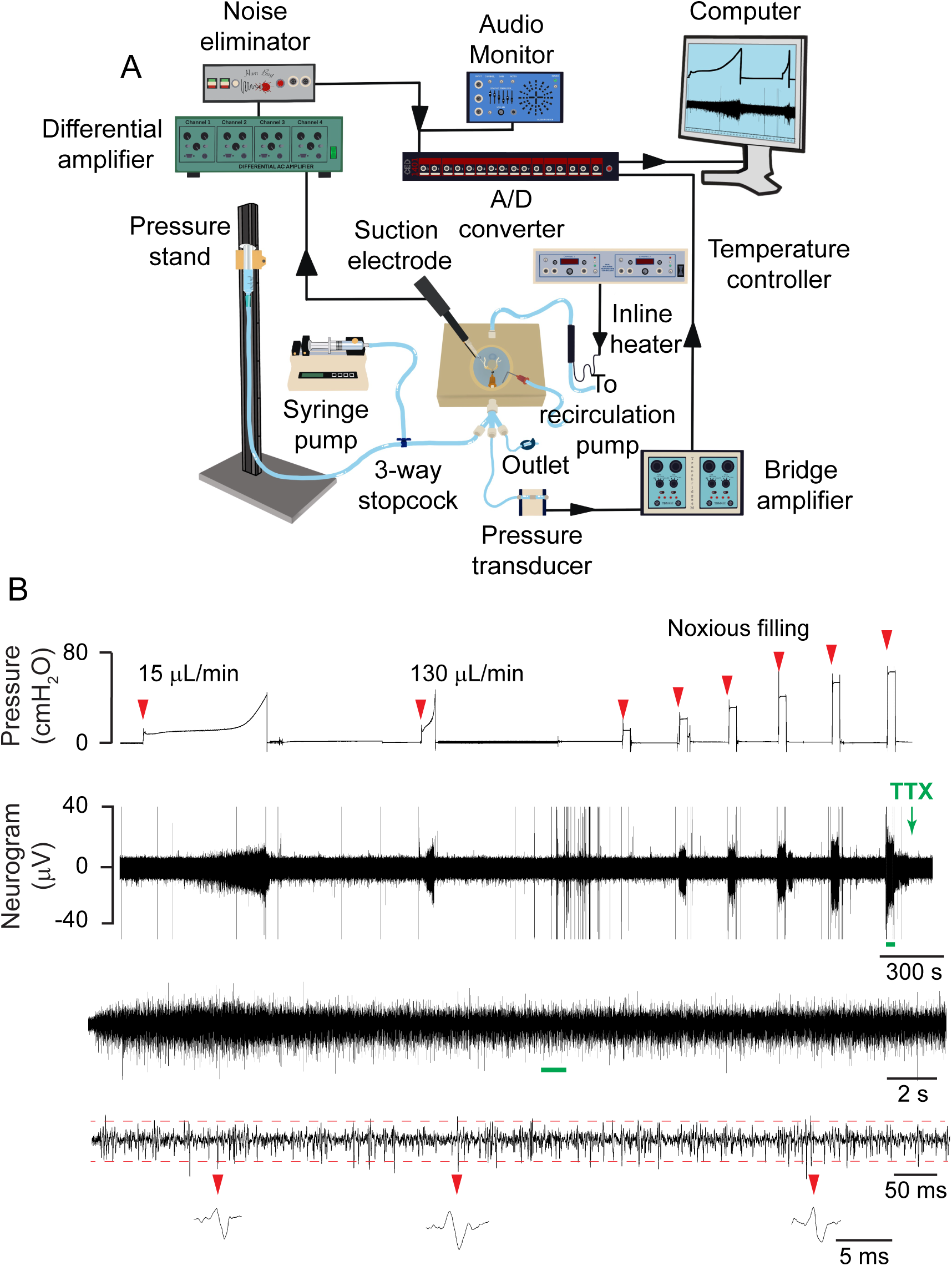
Ex vivo bladder-nerve preparation setup, action potential recording, and data analysis. (A) The urinary bladder and associated nerves were harvested, a cannula was inserted in the urethra and the ureters were tied close to the urinary bladder wall. The preparation was moved to a recording chamber containing warm Krebs solution continuously bubbled with 95% O₂/5% CO₂. The L6 or S1 spinal root was carefully placed onto a bipolar suction electrode to record afferent nerve activity. Electrical signals were differentially amplified, band-pass filtered at 100-1,000 Hz and then processed through a Hum Bug Noise Eliminator to remove 50/60 Hz noise and harmonics. The urethral cannula was coupled to a four-way connector: one branch led to a pressure transducer, a second port was connected to a syringe pump and a pressure stand for bladder filling, and the third port to a cannula with a pressure release valve for bladder emptying. (B) Representative multi-unit nerve recording. The urinary bladder was initially infused at a slow rate (15 µl/min) and subsequently at a fast rate (130 µl/min) up to a maximum pressure of 40 cmH₂O, followed by rapid at intravesical pressures of 10, 20, 30, 40, 50, and 60 cmH₂O. At the conclusion of each stimulus, the bladder was drained via the urethral cannula by manually opening the release valve. Red arrows indicate bladder filling. The neurogram was expanded (green line) to show individual spikes. The action potential detection threshold (red dotted lines) was set at three times the standard deviation of the baseline electrical signal recorded in the presence of tetrodotoxin (TTX). Individual spikes are displayed below the recording trace.

### Bladder afferent nerve recordings

To record afferent nerve activity, a spinal root (L6 or S1) was carefully positioned into a bipolar suction electrode (A-M system, Cat. No. 573040) suited with a glass pipette. The tip of the glass pipette was adjusted to fit the size required for whole-nerve or multi-unit recordings. Electrical signals were recorded with a differential amplifier Model 1700 (A-M Systems) and were band-pass filtered at 100-1,000 Hz. The amplified signal was further passed through a Hum Bug Noise Eliminator (Digitimer) to remove 50/60 Hz noise and harmonics. Pressure and nerve signals were recorded at a rate of 25,000 with a CED 1401 Power 3A data acquisition system (Cambridge Electronic Design) interfaced to a PC computer running Spike 2 software (Cambridge Electronic Design). For whole-nerve and multi-unit recordings, the preparation was equilibrated in the chamber with the bladder empty for at least 30 min. To assess the viability and reproducibility of the afferent response to filling, urinary bladders were continuously infused at a rate of 15 μl/min until the intravesical pressure reached 40 cmH_2_O (29 mmHg). This pressure was chosen because it reflects the peak pressure during cystometrograms in mice (38, 39). Once this pressure was reached, the bladder was emptied through the urethral cannula by manually opening the release valve. This protocol was repeated once more with a resting interval between fillings of 10 minutes.

### Experimental protocol for nerve recordings

To record afferent activity, bladders were filled at 15 μl/min until the intravesical pressure reached 40 cmH₂O (slow filling), and after a 10 min rest, intravesical pressure was increased to 10, 20, 30, 40, 50, and 60 cmH₂O for 30 s (noxious filling) with 3 min intervals between stimuli. To assess the effect of the filling rate on firing, an additional filling ramp at 130 μl/min to a pressure of 40 cmH_2_O was generated between the physiological filling and the noxious filling steps. Finally, for single-unit analysis, tetrodotoxin (TTX) at a final concentration of 1 μM was perfused into the chamber to block nerve activity and assess background noise.

### Nerve recordings data analysis

All nerve recordings were analyzed offline with Spike2. The threshold for detection of afferent activity for whole-nerve recordings was set at twice the root mean square (RMS) of the electrical signal when the bladder was empty (Fig. 3). The frequency of firing at a given pressure was calculated for an interval of 5 s or 0.58 s interval for bladder distended at 15 or 130 μl /min, respectively. Afferent activity was normalized to the firing frequency obtained at 40 cmH_2_O, setting this value as 100%. To determine the pressure required to increase afferent activity to 50% of the maximum, the relationship between pressure and normalized firing frequency was fitted to an allosteric sigmoidal equation, Ff= Ff_max_ × Ph / (P_50_h + Ph), where Ff is the normalized firing frequency at a given pressure, Ff_max_ is the maximum firing frequency, P is pressure, P_50_ is the pressure required to achieve 50% of the maximum afferent activity, and h is the Hill slope. The volume required to reach an intravesical pressure of 40 cmH_2_O was considered the maximal bladder capacity. Bladder capacity at a given pressure was calculated as the product of the infusion rate (15 µl/min or 130 µl/min) and the time required to reach a given pressure. Bladder capacity was normalized to the volume required to achieve an intravesical pressure of 40 cmH_2_O, setting this value as 100%. To determine the pressure required to achieve 50% bladder capacity, the relationship between intravesical pressure and normalized bladder capacity was fitted to an allosteric sigmoidal equation, Cap = Cap_max_ × Ph / (P_50_h + Ph), where Cap is the normalized bladder capacity at a given pressure, Cap_max_ is the maximum normalized bladder capacity, P is pressure, P_50_ is the pressure required to achieve 50% bladder capacity, and h is the Hill slope.

Single-unit analysis was conducted offline with Spike2 using spike detection and sorting via the New WaveMark function. To distinguish true neural spikes from noise background, a detection threshold was set at three times the standard deviation of the electrical signal in the presence of TTX. Before template matching, the DC offset of each snippet was removed to eliminate baseline fluctuations that do not reflect neuronal spikes and to ensure that template matching was based solely on waveform attributes relevant to individual fiber activity. Waveform snippets were extracted using the standard template settings, with cubic spline interpolation applied for waveform reconstruction. A 2.6 ms high-pass time constant was used to effectively isolate high-frequency signals and enhance spike discrimination. To ensure that most units were included in the analysis, the New WaveMark function was initially performed on a 60 s window of the recording corresponding to the last noxious stimulus (bladder distension at 60 cmH₂O for 30 s) and the 30 s preceding it. Spikes were manually reviewed and sorted as needed to ensure consistent unit discrimination. Once spikes were identified, the New WaveMark function was applied using the templates generated to the rest of the recording.

### Muscle strips contractility measurements

Following euthanasia, an abdominal vertical incision was performed, and the bladder was exposed and removed by cutting it at the neck, close to the proximal urethra. The tissue was placed in a Sylgard coated dish filled with room temperature Krebs solution (composition in mM: NaCl 118, KCl 4.7, CaCl_2_ 1.9, MgSO_4_ 1.2, NaHCO_3_ 24.9, KH_2_PO_4_ 1.2, glucose 11.7) bubbled with a mixture of 95% O_2_/5% CO_2_ (v/v). Bladders were cut open in half and pinned flat, and silk sutures were used to secure both ends of the bladder strips. The strips were transferred to the experimental chambers where they were completely immersed in 15 ml of bubbled Krebs solution and kept at 37°C via a circulating water pump. One end of the strip was attached to a fixed glass rod, and the other to a force transducer, to measure tissue contraction. Tissue was stretched by applying 10 mN (1 g) baseline tension and allowed to equilibrated in the chambers for 30 min (40). Electrical field stimulation (EFS) was applied with an S88 stimulator (Grass Instruments) and delivered in 5 s pulses using two platinum electrodes placed on each side of the strips. Sequential stimulation pulses with an intensity of 80 V and frequencies of 1, 2.5, 5, 7.5, 10, 12.5, 15, 20, 30, 40, and 50 Hz were applied to generate EFS frequency-response curves. After the last stimulus applied, DMSO (vehicle, 15 µl) or Yoda1 (30 µM) were added directly to the bath, and another frequency-response curve was generated. Then, 1 µM TTX was added to the bath to verify that EFS-evoked smooth muscle contractions were driven by nerve transmission. KCl (80 mM) was added at the end of the experiments to confirm tissue viability (40, 41).

### Muscle strips data analysis

Data were acquired using WinDaq software (DATAQ Instruments) and analyzed with Spike2. Amplitude (g) of EFS-induced responses was calculated and normalized to the maximal contraction (50 Hz, highest frequency applied) obtained from the control. When two strips from the same mouse received the same treatment, their values were averaged. To assess changes in basal smooth muscle activity, the RMS of the signal was measured for a 5 min interval before and after the addition of Yoda1 or DMSO.

### Retrograde labeling of bladder sensory neurons

Bladder dorsal root ganglia (DRG) neurons were labeled with unconjugated cholera toxin β subunit (CTb) (Sigma, Cat. N°. C9903). Mice were anesthetized with isoflurane and an abdominal incision was made to expose the bladder. Then, 2 μl of CTb (0.1% in sterile saline) was injected at three to four locations in the bladder wall with a Hamilton^TM^ 600 Microliter syringe (Hamilton, Cat. N°. 763301) suited with a 33G needle (Hamilton, Cat. N°. 7803-15). After each injection, the needle was kept in place at the site for 20-30 s. The muscle layer and skin incisions were closed in layers using a 5.0 PDO absorbable monofilament surgical suture (AD Surgical, Cat. N°. S-D518R13). Mice were administered ketoprofen (5 mg/kg, Zoetis, Ketofen) subcutaneously for pain relief, and ampicillin (100 mg/kg, Eugia US LLC, Cat. No. NDC 55150-113-10) to prevent postoperative infections. Mice were housed under the conditions described above for 7 days before DRG harvesting.

### Immuno-fluorescence in situ hybridization (Immuno-FISH), including image capture

The RNAscope^TM^ Multiplex Fluorescent v2 kit (ACD, Cat. N°. 323100) was used to examine expression of *Piezo1* and *Calca* in DRG. The following probes were used: *Piezo1* (Cat. N°. 400181-C1) and *Calca* (Cat. N°. 578771-C2). A negative control probe (ACD, Cat. N°. 320871) was used to detect non-specific signals. L6-S2 DRG were harvested after euthanasia. Tissues were dipped into a dish with Optimal Cutting Temperature compound (OCT; Fisher Scientific, Cat. N°. 23-730-571) and then transferred to a 10 x 10 x 5 mm Tissue-Tek Cryomold (Sakura Finetek USA Inc, Cat. N°. 427971) filled with OCT that was later placed in a -80 °C freezer. Sections (12 μm) were cut with a Leica CM1950 cryostat (chamber temperature of -20 °C and a knife temperature of -18 °C) and collected on Superfrost^TM^ Plus microscope slides (ThermoFisher Scientific, Cat. N°. 12-550-15). Slices containing sectioned tissues were allowed to “dry” in the cryostat chamber for 30 min, before long-term storage at -80 °C. Slides containing sectioned tissue were removed from the -80 °C freezer and immediately fixed for 15 min at 4 °C with neutral buffered formalin comprised of 29 mM NaH_2_PO_4_•H_2_O, 45.8 mM Na_2_HPO_4_, and 4.0% (v/v) paraformaldehyde. Sections were incubated overnight with a rabbit antibody against CTb (Novus Biologicals, Cat. N°. NB100-63067, dilution 1:100) at 4 °C. After primary antibody incubation, slides were incubated in neutral buffered formalin for 30 min at room temperature, and then with Protease IV for 5 min at room temperature. Probes were developed using TSA Vivid^TM^ Fluorophore 520 (TocrisBiosciences, Cat. N°. 7523), TSA Vivid^TM^ Fluorophore 570 (Tocris Biosciences, Cat. N°. 7526). Following the last treatment with horseradish peroxidase blocker, samples were incubated for 30 min at room temperature with a donkey anti-rabbit secondary antibody conjugated with AlexaFluor^TM^ 647 (1:100). Nuclei were counterstained with DAPI. Tissues sections were mounted using ProLong Gold antifade mountant (ThermoFisher Scientific, Cat. N°. P36934) and cured overnight at room temperature in the dark. The slides were then stored at 4 °C. General image capture was performed using a Leica HCX PL APO 20x, 0.75 numerical aperture dry objective and the appropriate laser lines of a Leica Microsystems SP8 Stellaris confocal microscope outfitted with a 405-laser diode and a white light laser. 8-bit images were collected at 600Hz using 2-line averages combined with 2 frame averages. Sequential scanning was used to prevent cross talk between channels. Stacks of images (1,024 x 1,024, 8-bit) were collected using system-optimized parameters for the z-axis. Images were processed using the 3D visualization package of Leica LASX software and exported as TIFF files. Final figures were assembled in Adobe Illustrator version 29.6.1 (Adobe).

### Statistical analysis

Data are expressed as mean ± SEM (n), where n equals the number of independent measurements. Parametric or nonparametric tests were employed as appropriate. A p value < 0.05 was considered statistically significant. Statistical comparisons were performed with GraphPad version 11.0.1 (GraphPad Software, San Diego, CA).

## Results

### Functional classification of bladder afferents

To investigate the response of bladder afferents to physiological, fast and noxious distension, we used an *ex vivo* bladder-pelvic nerve preparation and recorded simultaneously intravesical pressure and action potentials evoked by filling (Fig. 1A). Single units were isolated from multi-unit recordings and sorted offline using Spike2. In preliminary experiments, we noticed the difficulty in classifying fibers based on their activation threshold when the urinary bladder was filled at a physiological rate (15 μl/min). For instance, some fibers show a rapid increase in firing rate as the bladder fills, whereas others discharge action potentials sporadically before suddenly exhibiting a sharp rise in firing rate. In addition, some fibers begin firing occasional action potentials at intravesical pressures above 10 cmH₂O, but their firing rate never increased beyond that. In our studies, the pressure at which the units begin to discharge does not correlate well with their behavior as the bladder fills. Thus, to classify mechanosensitive afferents we developed a two-step filling protocol: the initial filling phase was at a physiological rate (15 μl/min), and the second phase involved noxious 30 s fast filling steps with 3 min intervals between them. The later step protocol has been broadly used to trigger a visceromotor response (VMR), a reflex contraction of the abdominal musculature evoked in response to noxious stimulation of the bladder or colon (42, 43). At the end of the experiment the chamber was perfused with tetrodotoxin (TTX) to block nerve activity and determine the baseline. A representative recording illustrating the setup, stimulation protocol and the approach for sorting is shown in Fig. 1. Firing rates for individual units were calculated for both filling protocols (i.e., physiological and noxious) as described in Materials and Methods.

Visual inspection of firing plots of individual units reflects the presence of three distinctive types: 1) a group that responds to physiological filling, with its firing rate increasing linearly with intravesical pressure (type I), 2) a second group that also responds to physiological filling, but whose firing plateaus as the intravesical pressure rises (type II), and 3) a third population of fibers that is near silent during physiological filling and is activated by noxious pressures (type III) (Fig. 2A). A fourth class of afferents that exhibits constitutive firing and is inhibited by noxious stimuli was occasionally observed. Due to the low frequency of appearance, this type of afferent was not further studied. Type I and II afferents have similar threshold of activation but differ in the response to distension during physiological and noxious filling. To differentiate and adequately classify type I and II fibers, we developed an unbiased approach that models the relationship between intravesical pressure (10-40 cmH2O) and firing rate using linear regression. If the slope was > 0.1 and significantly different from zero, then units were considered type I. If the slope of the pressure-firing rate relationship was < 0.1 or nonlinear, units were counted as Type II. Type I fiber firing increases linearly as the bladder fills, up to pressures of 40 cmH_2_O (86/243, 35.4%). Type II afferent firing increases suddenly and plateaus at pressures < 20 cmH_2_O (61/243, 25.1%). Furthermore, Type I and II afferents exhibit different responses to noxious filling, with type I firing plateauing at 40 cmH_2_O and type II firing increasing linearly up to 60 cmH_2_O. In contrast, Type III afferents are nearly silent during physiological filling, but they respond to sudden increases in intravesical pressures (95/243, 39.5%). Representative firing of individual units from an experiment before and after sorting using the approach described in Materials and Methods is shown (Fig. 2B-C). Data from 243 fibers responding to physiological and noxious filling are presented in Fig. 2D-F. In summary, our findings reveal the existence of three distinct subsets of bladder afferents that are activated by physiological and noxious distension of the urinary bladder.

**Figure 2.**
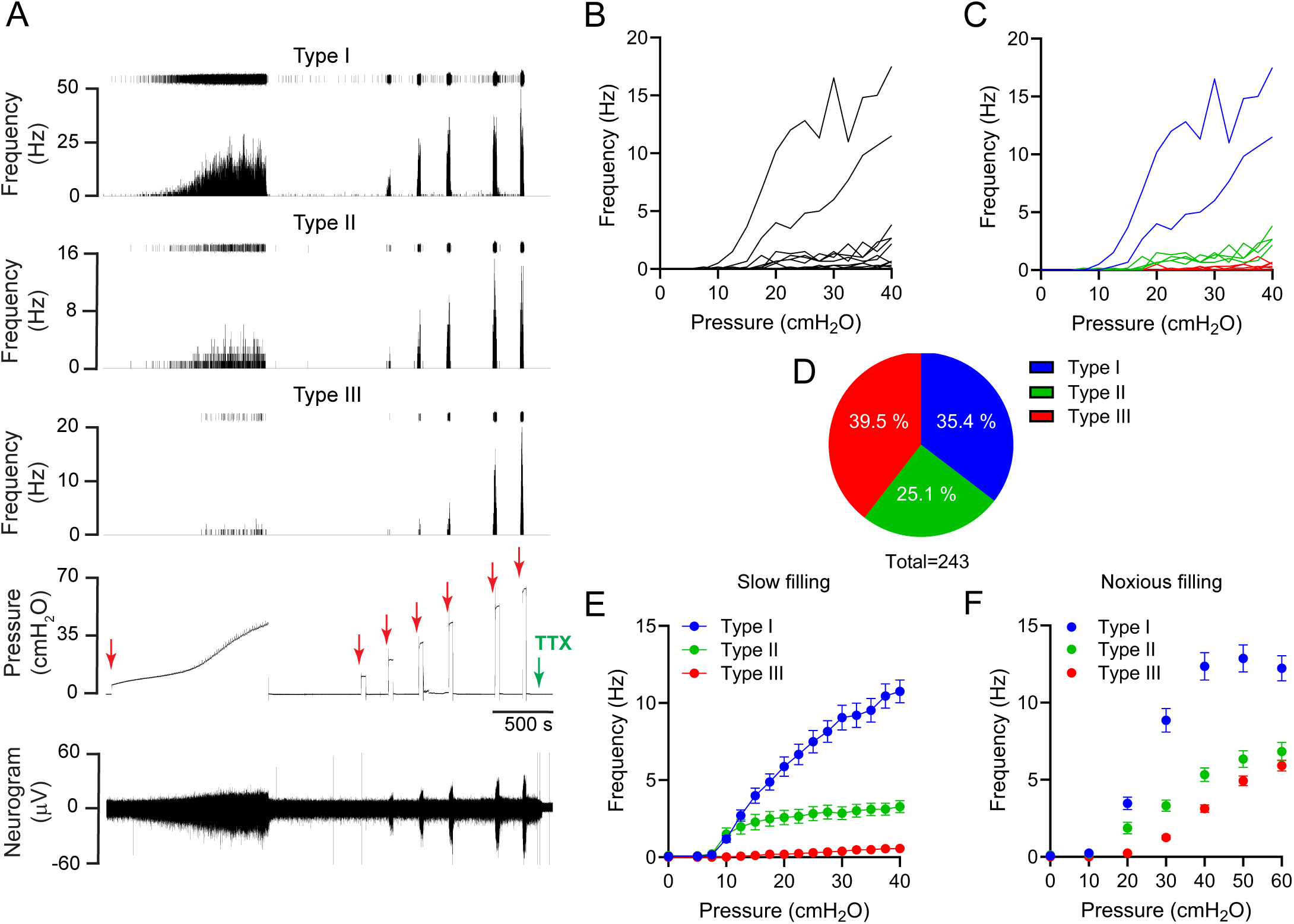
Classification of mechanosensitive fibers innervating the urinary bladder. (A) Representative extracellular recording illustrating the activity of the three principal types of mechanosensitive afferent fibers. The urinary bladder was filled at 15 µl/min up to 40 cmH₂O (physiological filling), and then rapidly filled for 30 s at 10, 20, 30, 40, 50, and 60 cmH₂O (noxious filling). A representative example of each type of fiber is displayed. For each fiber, the instantaneous firing rate (Hz) and individual spikes (top) are displayed. Red arrows indicate bladder filling. (B, C) Firing rates of single units before (black) and after (colored) spike sorting. Units were classified according to their response profile during physiological and noxious filling as described in Materials and Methods. Type I afferents are shown in blue, type II in green and type III in red. (D) Distribution of mechanosensitive bladder afferent fiber types. A total of 243 single units were recorded and classified. (E and F) Discharge pattern of each afferent type during slow (E) or noxious (F) bladder distension. A total of 243 single units were recorded and classified (type I 86 units, type II 61 units, and type III 96 units). Data are shown as the mean ± SEM (n=21; the responses to slow and noxious filling differed significantly across afferent types, two-way ANOVA, **** p<0.0001).

### Filling rate differentially affects firing of bladder afferent types

To assess whether the three identified afferent populations represent different entities, we investigated the effect of fast bladder filling on firing. Prior work has established that the bladder filling rate directly influences afferent function, with faster filling rates associated with greater afferent activity (44). Initially, we conducted whole-nerve recordings to assess the effect of fast filling on afferent discharge and bladder wall mechanics and elasticity. Urinary bladders were initially infused at a rate of 15 μl/min until the intravesical pressure reached approximately 40 cmH_2_O, and subsequently at a rate of 130 μl/min to the same final pressure (Fig. 3A). At a filling rate of 15 μl/min, afferents begin to discharge above basal levels when the intravesical pressure reaches approximately 7-8 cmH_2_O (Fig. 3A-B). Fast filling shifts the pressure at which afferents begin to discharge to approximately 10 cmH_2_O. The pressure required to increase afferent activity to 50% of the maximum was 20.94 cmH₂O (95% CI, 20.05-21.83) at a filling rate of 15 μl/min and 24.95 cmH₂O (95% CI, 24.12-25.78) at a filling rate of 130 μl/min. Likewise, the relationship between bladder capacity and intravesical pressure was shifted to the right with fast filling. The pressure required to achieve 50% bladder capacity was 12.25 cmH₂O (95% CI, 11.62-12.97) at a filling rate of 15 μl/min and 18.83 cmH₂O (95% CI, 17.98-19.89) at a filling rate of 130 μl/min. These findings suggest that afferent activity depends, at least in part, on the extent of bladder filling. However, the apparent bladder capacity, which represents the volume required to increase the intravesical pressure to 40 cmH_2_O, was similar at filling rates of 15 or 130 μl/min (Fig. 3D). Overall, our studies suggest that the filling rate influences bladder wall mechanics and elasticity, and indirectly afferent firing.

**Figure 3.**
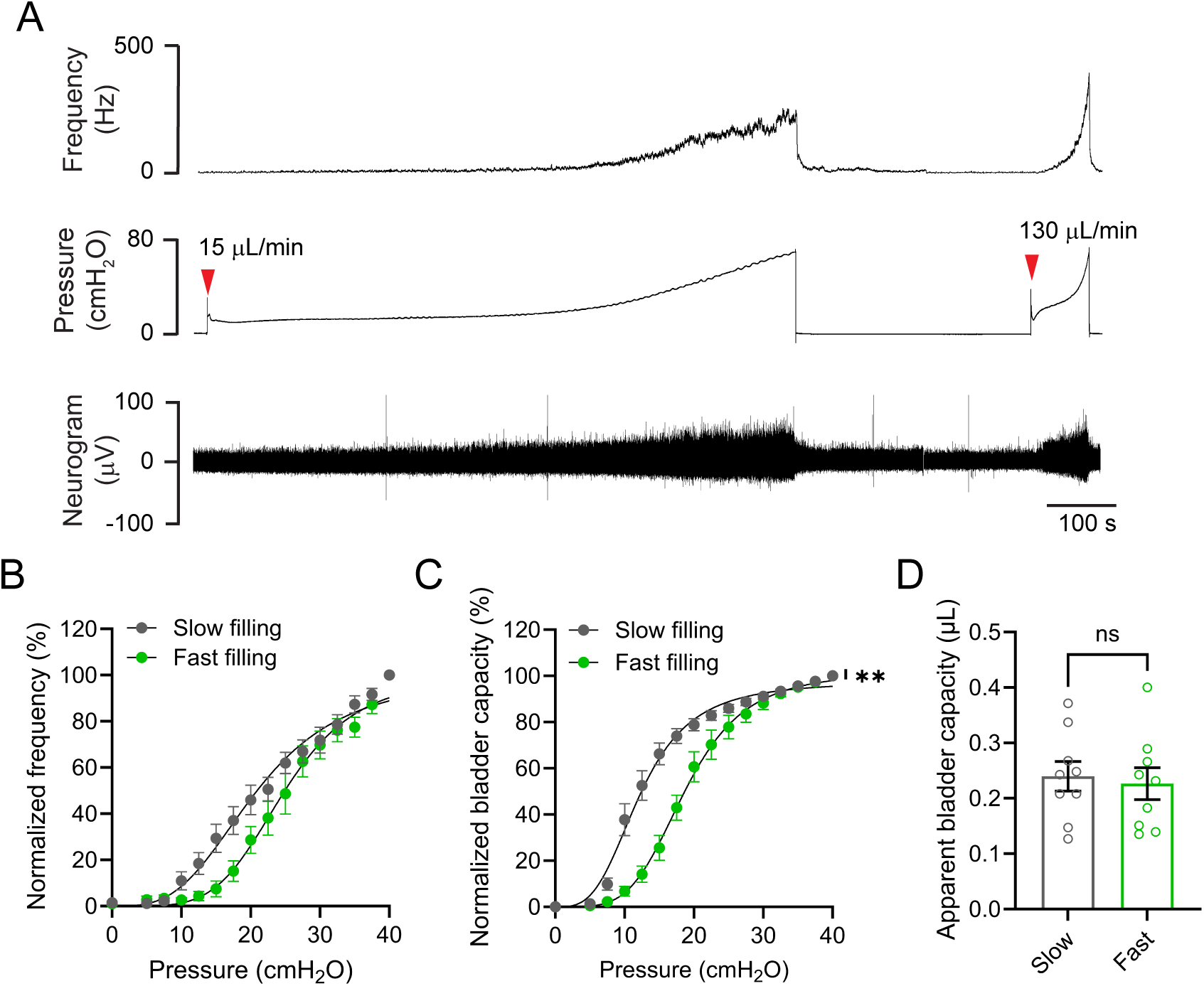
Filling rate influences afferent discharge. (A) Representative whole-nerve recording illustrating the effect of the filling rate on afferent activity. The bladder was continuously filled first at a slow rate of 15 µl/min and subsequently at a faster rate of 130 µl/min, up to a maximum pressure of 40 cmH₂O. Red arrows indicate bladder filling. (B and C) Normalized afferent firing (C) and normalized bladder capacity (C) as a function of the intravesical pressure. Data are shown as the mean ± SEM (n=9; two-way ANOVA, * p<0.01). (D) Apparent bladder capacity for slow filling and fast filling. Apparent bladder capacity represents the volume required to increase the intravesical pressure to 40 cmH_2_O.

We next performed single-unit analysis to investigate the responses to slow and fast filling of the different types of bladder afferents. In these experiments, bladders were filled at 15 μl/min and subsequently at 130 μl/min (each to 40 cmH₂O), followed by a series of 30 s fast-filling steps to pressures ranging from 10 to 60 cmH₂O. Mechanosensitive afferents were classified using the criteria described above. As seen in Fig. 4, fast filling increases the maximal firing rate of the three types of afferent fibers, but it does not appear to change the threshold pressure for activation (Fig. 4 A-C). Thus, the increase in firing observed with fast filling occurs once the activation threshold is surpassed. These findings indicate that the rate of bladder filling controls afferent firing discharge but not the pressure at which afferents are recruited.

**Figure 4.**
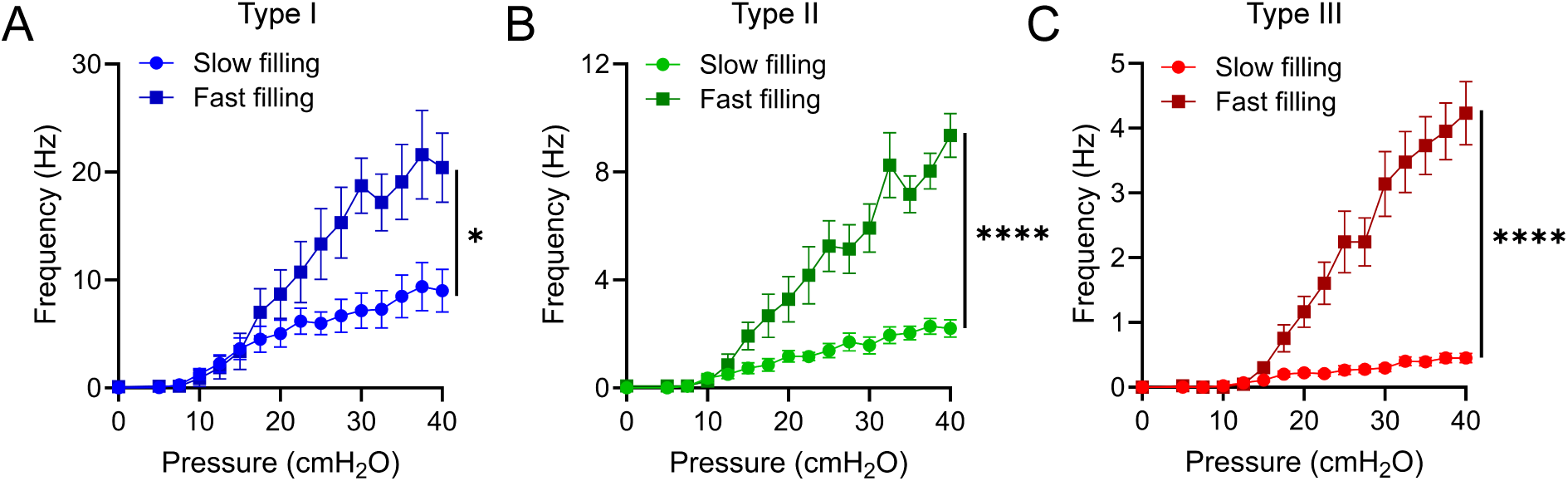
Fast bladder filling differentially modulates the firing pattern of mechanosensitive afferent fiber types. (A, B and C) Discharge pattern of afferent fiber type I (A), II (B), and III (C) during slow (15 µl/min) and fast (130 µl/min) filling. A total of 59 single units were recorded and classified. Data are shown as the mean ± SEM (n=7; two-way ANOVA, * p<0.05 and **** p<0.0001).

### Yoda1 regulates afferent activity in a fiber-type-specific manner

Piezo channels are trimeric channels that sense mechanical forces in a variety of tissues, playing a crucial role in a broad range of mechanotransduction processes. In the urinary bladder, PIEZO1 channels are expressed in the urothelium, smooth muscle cells, endothelial cells and fibroblasts (45). PIEZO2 channels are primarily expressed in bladder sensory neurons and umbrella cells (46). Both PIEZO1/2 in urothelial cells and PIEZO2 in sensory neurons contribute to mechanotransduction in the urinary bladder, thereby participating in the regulation of voiding (41, 46). We used Yoda1, a specific activator of PIEZO1, to examine the effect of PIEZO1 activation on afferent function. Yoda1 specificity toward PIEZO1 has been validated in multiple studies using knockout mice (41, 47–52). Thus, bladders were slowly filled at 15 μl/min until the pressure reached 40 cmH_2_O and subsequently by 30 s fast filling steps (to 10, 20, 30, 40, 50 or 60 cmH_2_O) with 3 min intervals between them. This protocol was repeated after the addition of 30 μM Yoda1 to the bath. Yoda1 caused a slightly, but significant, increase in the basal activity of type I and II afferents (Fig. 5 G-H), but it did not shift the threshold pressure for activation of any bladder afferent subtype (Fig. 5 A-C). Yoda1 significantly increased the rate of firing of type I and III afferent during slow filling (Fig. 5 A-C), and of type III during noxious stimulation (Fig. 5 D-F). The vehicle control (DMSO) did not significantly alter afferent discharge. These results indicate that Yoda1 modulates bladder afferent activity in a fiber-type-specific manner. Taken together, our studies suggest that PIEZO1 regulates the intensity of the response to distension of type I and III afferents rather than determining when afferents are recruited.

**Figure 5.**
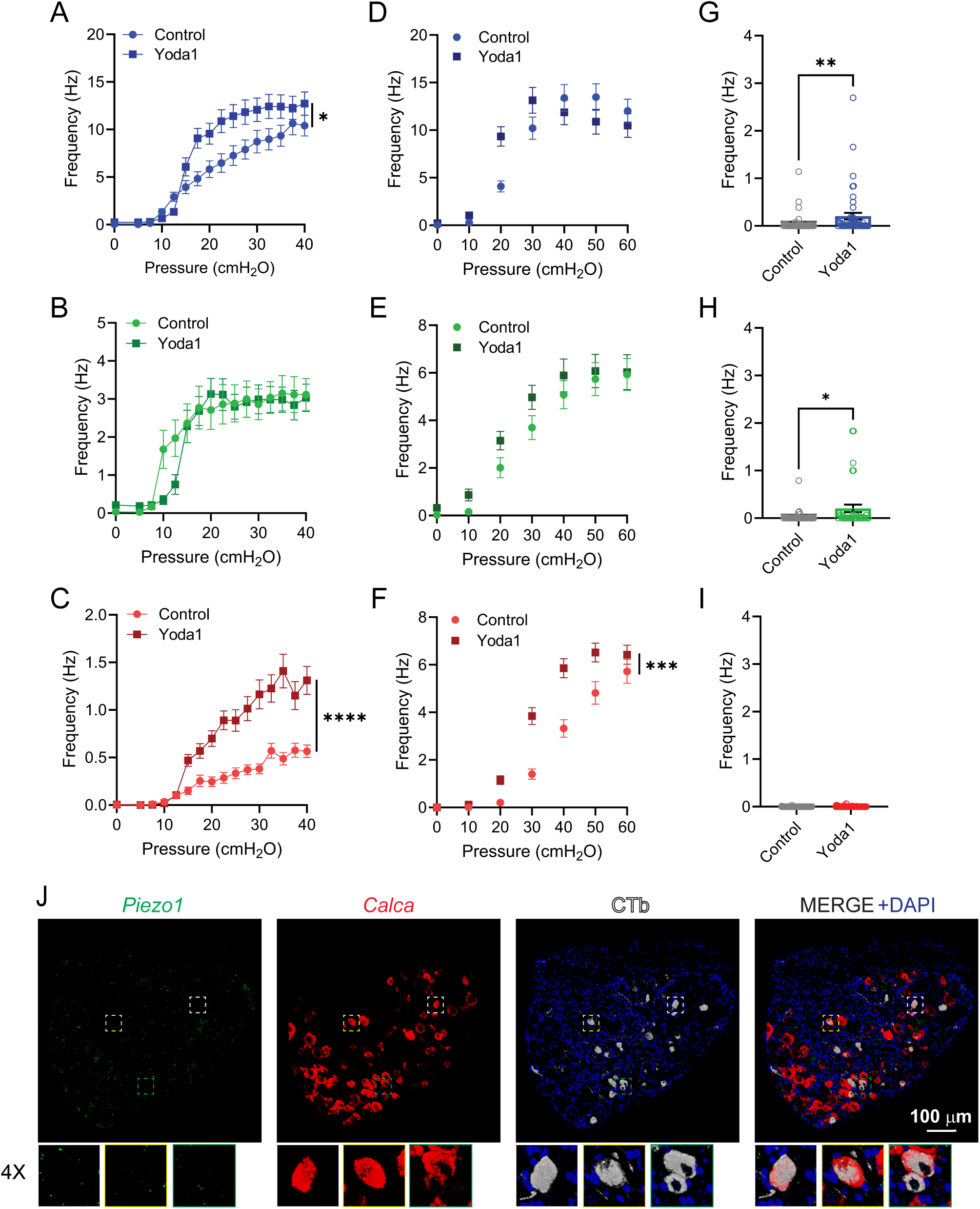
Differential regulation of mechanosensitive bladder afferent types by Yoda1. (A, B, C) Discharge pattern of mechanosensitive afferent fiber types I (A), II (B), and III (C) during slow bladder filling in the absence and presence of Yoda1 (30 µM). (D, E, F) Discharge pattern of mechanosensitive afferent fiber types I (D), II (E), and III (F) during noxious filling in the absence and presence of Yoda1 (30 µM). (G, H, I) Basal afferent activity of mechanosensitive afferent fiber types I (G), II (H), and III (I). A total of 137 single units were recorded and classified. Data are shown as the mean ± SEM (n=11; two-way ANOVA, * p<0.05, ** p<0.01 and **** p<0.0001). (J) Immuno-FISH confirms negligible expression of *Piezo1* in bladder sensory neurons. No signal was visible with the negative control probe (not shown). Confocal images of L6-S2 dorsal root ganglia (DRG) are shown. Immuno-FISH was performed in fresh frozen sections of DRG harvested from mice injected into the bladder wall with cholera toxin β subunit (CTb). A rabbit antibody anti-CTb and a donkey anti-rabbit secondary antibody conjugated with AlexaFluor^TM^ 647 were used to identify bladder sensory neurons (white). Inset, 4-fold magnification of CTb-labeled neurons. Representative of 3 independent experiments (n=6 mice).

Previous studies have shown that Piezo1 channels are expressed in a subset of sensory neurons, where they contribute to the sensation of mechanical itch (53). To establish whether direct activation of Piezo1 channels within type I and III afferent fibers underlies the Yoda1-evoked increase in firing, we combined immuno-FISH with retrograde tracing using CTb (Fig. 5J). *Calca*, a marker of peptidergic sensory neurons, was used as positive controls in these studies. As shown in Fig. 5J, intracellular mRNA clusters are present in some bladder-innervating sensory neurons, though overall expression is low. Many cells do not express *Piezo1*. These findings suggest that the increase in afferent discharge observed with Yoda1 cannot be attributed to direct activation of PIEZO1 channels in afferents.

### Yoda1 does not alter smooth muscle contractibility

Previous studies have shown that transient contractions of the urinary bladder smooth muscle promote afferent activity during filling (54). To determine whether Yoda1 increases afferent firing by activating PIEZO1 channels in smooth muscle cells, we compared nerve-induced detrusor contractions in the absence and presence of Yoda1 (30 μM). EFS stimulation parameters including frequency, duration and voltage were based on previous studies (40, 55, 56). Stimulus-response curves for bladder strips in the absence and presence of Yoda1 or DMSO (vehicle) are shown in Fig. 6. Yoda1 did not affect basal activity or EFS-induced detrusor contractions. As expected, TTX blocked nerve-mediated smooth muscle contractions. Thus, these findings suggest that the Yoda1-induced increase in afferent activity is not mediated by smooth muscle activation.

**Figure 6.**
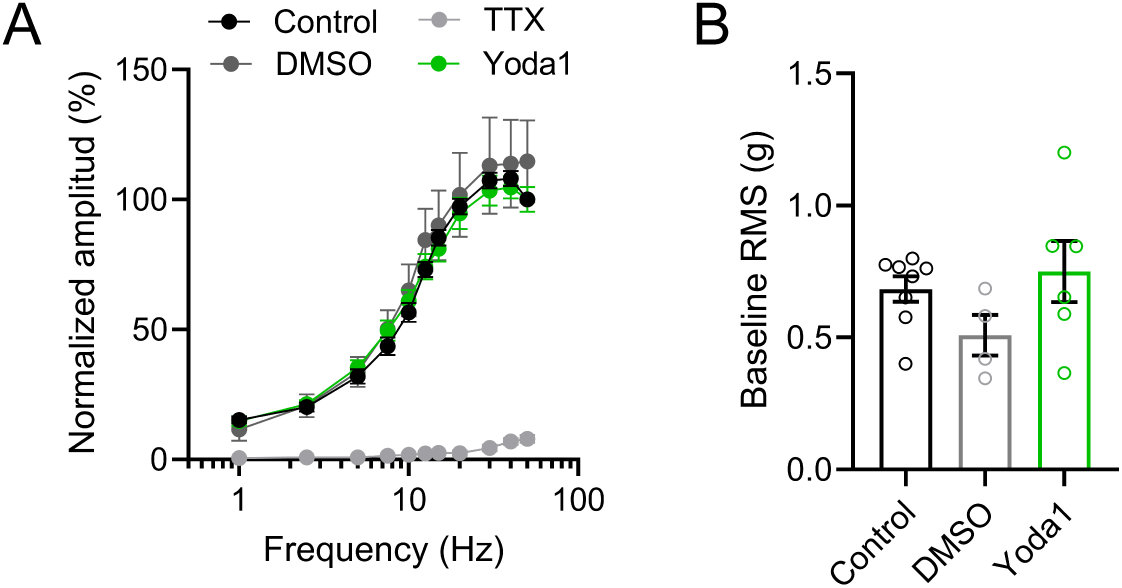
Yoda1 does not alter smooth muscle contractility. Electric field stimulation (EFS) of mouse bladder urothelium-intact muscle strips. (A) Stimulus-response curves elicited by EFS before (control) and after DMSO (vehicle), Yoda1 (30 µM) and tetrodotoxin (TTX, 1 µM). Yoda1 (n=6) or DMSO (n=4) do not alter the response of the bladder smooth muscle to EFS. TTX was added at the end of each experiment to verify that EFS-evoked smooth muscle contractions were driven by nerve transmission. (B) Effect of Yoda1 or vehicle on basal contractile activity.

## Discussion

We developed a systematic two-step approach to classify functionally bladder afferents based on their response to physiological and noxious mechanical stimulation. To identify single units and generate templates for spike discrimination, we analyzed responses to noxious mechanical stimulation at 60 cmH₂O. These templates were then used to analyze afferent responses to slow, fast, and noxious filling. Three major populations of bladder afferents were identified: two activated during physiological distension and one recruited by noxious mechanical stimulation. This latter population would have been impossible to detect without the two-step approach described here. Consistent with our findings, Mills et al. recently reported two types of low-threshold afferents innervating the mouse urinary bladder: wide-range afferents, whose firing increases exponentially with pressure, and narrow-range afferents, whose firing plateaus at approximately 20 cmH₂O (57). A third type of afferent with a higher threshold (7-15 cmH_2_O) and firing that increases almost linearly with pressure was also described (57). Although the response to bladder filling of these high-threshold afferents differs from that of the type III afferents described in our report, it is possible that they represent the same population and that their distinct behavior is simply a consequence of the experimental conditions. Our findings do not rule out the existence of additional afferent populations that are silent under the conditions used in this study but may become mechanosensitive upon appropriate stimulation.

The threshold pressure of activation has been broadly used to classify bladder afferents, with 20 cmH₂O being the cutoff chosen because it is close to the micturition threshold in rodents (10, 22, 37, 57). We found that the threshold was not a reliable way to classify bladder afferents when the bladder is filled at near physiological rates (15 μl/min). This led us to develop a new approach to classify bladder afferents. To assess whether the populations identified in our study were indeed different entities, we examine the response to fast filling and to the PIEZO1 channel activator Yoda1. Fast filling did not change the activation threshold of the three types of afferents, but it increased their rate of discharge in a type-specific manner. The rate of firing at 40 cmH₂O increased approximately twofold for type I afferents, fourfold for type II afferents, and tenfold for type III afferents. Likewise, Yoda1 increased the rate of discharge of type I and III afferents during slow filling, and it shifted the activation curve for type III increasing their sensitivity to noxious filling. In addition, Yoda1 slightly increased basal firing of type I and II afferents. Taken together, these findings indicate that type I, II, and III afferents represent functionally distinct populations with unique mechanosensory properties, and that their classification based on response patterns to multiple stimuli is more reliable than threshold-based classification alone.

Bladder sensory neurons play a critical role in detecting distension and noxious mechanical stimuli, yet the mechanisms underlying these processes remain poorly understood. Unlike mechanotransduction in cutaneous sensory organs and proprioceptors, which occurs within milliseconds (58, 59), filling of the bladder in mammals takes minutes to hours. Furthermore, while skin and skeletal muscle afferents are associated with specialized sensory organs, bladder afferents have free nerve terminal endings (58), suggesting that other structures must contribute to mechanosensation in the bladder wall. Bladder filling initiates a series of morphological changes, including thinning of the umbrella cell layer and lamina propria, unfolding of the mucosal rugae, and stretching of the smooth muscle. As afferents are distributed throughout the mucosa and muscularis externa, any of these structural changes could potentially serve as a stimulus for afferent activation. For instance, purinergic signaling through P2rx2 and P2rx3, initiated by stretching of bladder tissues, was proposed to be involved in mechanosensory transduction in the urinary bladder (60–62). Work by Rong et al. provided support to the notion that ATP released endogenously in the bladder contributes to sensory signaling across the full range of stimuli, from innocuous to nociceptive (63). However, it should be noted that, using the same mouse lines (61, 62), Takezawa et al. found no voiding defects in P2rx2 or P2rx3 knockout mice under baseline conditions, with alterations observed only following bacterial lipopolysaccharide administration (64). Other hypotheses have been proposed to explain the mechanisms underlying bladder afferent activation during filling (54, 65), but none have been conclusively demonstrated. In summary, our findings suggest that each of the bladder afferent populations identified here is governed by distinct activation and regulatory mechanisms.

PIEZO1 channels are expressed throughout multiple layers of the bladder wall, including the urothelium, PDGFRα+ fibroblasts of the lamina propria and detrusor, smooth muscle cells, and mesothelial cells (45). To assess whether PIEZO1 channels regulate the response of bladder afferents to distension, we compared their firing in the absence and presence of Yoda1. As described above, Yoda1 increases the firing rate of type I and II afferents during physiological filling and enhanced the sensitivity of type III afferents to noxious stimulation. Although PIEZO1 channels are expressed in smooth muscle cells and fibroblasts in the detrusor, EFS-evoked contractions in the presence of Yoda1 were not different from those of controls. These finding suggests that the effects of Yoda1 on afferent function are not attributable to changes in smooth muscle contractility. Likewise, although a contribution of PIEZO1 in afferents to the Yoda1-evoked increase in firing cannot be definitively excluded, immuno-FISH studies suggest its expression sensory neurons is negligible under basal conditions. We previously showed that Yoda1 causes a PIEZO1-dependent increase in intracellular Ca^2+^ in urothelial cells (41). Thus, one potential mechanism by which Yoda1 could increase afferent firing may involve the release of neuroactive substances from the urothelium or fibroblasts in the lamina propria.

In summary, our study provides a systematic approach for the functional classification of bladder and other visceral afferents, and provides a framework for understanding the function, regulation and potentially the relationship between their physiological response and molecular identity.

## Author contributions

MDC and GMM conceived and designed the study. GMM performed bladder afferent nerve recordings. GMM, SLD and JMB performed muscle strip studies. XS performed fluorescence *in situ* hybridization (FISH) staining. GMM and MDC analyzed data. GMM and MDC drafted the manuscript with input from all authors. MDC, GMM, SLD, JMB and XS edited and revised the manuscript. All the authors approved the final version of the manuscript.

## References

1. de Groat WC, Griffiths D, and Yoshimura N. Neural control of the lower urinary tract. Comprehensive Physiology 5: 327, 2015.

2. Vizzard MA, Erickson VL, Card JP, Roppolo JR, and de Groat WC. Transneuronal labeling of neurons in the adult rat brainstem and spinal cord after injection of pseudorabies virus into the urethra. J Comp Neurol 355: 629–640, 1995.

3. Nadelhaft I, Vera PL, Card JP, and Miselis RR. Central nervous system neurons labelled following the injection of pseudorabies virus into the rat urinary bladder. Neurosci Lett 143: 271–274, 1992.

4. Bahns E, Halsband U, and Janig W. Responses of sacral visceral afferents from the lower urinary tract, colon and anus to mechanical stimulation. Pflugers Arch 410: 296–303, 1987.

5. Habler HJ, Janig W, and Koltzenburg M. Myelinated primary afferents of the sacral spinal cord responding to slow filling and distension of the cat urinary bladder. J Physiol 463: 449–460, 1993.

6. Janig W, and Morrison JF. Functional properties of spinal visceral afferents supplying abdominal and pelvic organs, with special emphasis on visceral nociception. Progress in brain research 67: 87–114, 1986.

7. Bahns E, Ernsberger U, Janig W, and Nelke A. Functional characteristics of lumbar visceral afferent fibres from the urinary bladder and the urethra in the cat. Pflugers Arch 407: 510–518, 1986.

8. Floyd K, Hick VE, and Morrison JF. Mechanosensitive afferent units in the hypogastric nerve of the cat. J Physiol 259: 457–471, 1976.

9. Dmitrieva N, and McMahon SB. Sensitisation of visceral afferents by nerve growth factor in the adult rat. Pain 66: 87–97, 1996.

10. Sengupta JN, and Gebhart GF. Mechanosensitive properties of pelvic nerve afferent fibers innervating the urinary bladder of the rat. J Neurophysiol 72: 2420–2430, 1994.

11. Wen J, and Morrison JF. The effects of high urinary potassium concentration on pelvic nerve mechanoreceptors and ’silent’ afferents from the rat bladder. Adv Exp Med Biol 385: 237–239, 1995.

12. Sengupta JN, and Gebhart GF. Characterization of mechanosensitive pelvic nerve afferent fibers innervating the colon of the rat. J Neurophysiol 71: 2046–2060, 1994.

13. Xu L, and Gebhart GF. Characterization of mouse lumbar splanchnic and pelvic nerve urinary bladder mechanosensory afferents. J Neurophysiol 99: 244–253, 2008.

14. Zagorodnyuk VP, Brookes SJ, Spencer NJ, and Gregory S. Mechanotransduction and chemosensitivity of two major classes of bladder afferents with endings in the vicinity to the urothelium. J Physiol 587: 3523–3538, 2009.

15. Zagorodnyuk VP, Gibbins IL, Costa M, Brookes SJ, and Gregory SJ. Properties of the major classes of mechanoreceptors in the guinea pig bladder. J Physiol 585: 147–163, 2007.

16. Kuga N, Tanioka A, Hagihara K, and Kawai T. Fiber type-specific afferent nerve activity induced by transient contractions of rat bladder smooth muscle in pathological states. PloS one 12: e0189941, 2017.

17. Zagorodnyuk VP, Keightley LJ, Brookes SJH, Spencer NJ, Costa M, and Nicholas SJ. Functional changes in low- and high-threshold afferents in obstruction-induced bladder overactivity. Am J Physiol Renal Physiol 316: F1103–f1113, 2019.

18. McCarthy CJ, Zabbarova IV, Brumovsky PR, Roppolo JR, Gebhart GF, and Kanai AJ. Spontaneous contractions evoke afferent nerve firing in mouse bladders with detrusor overactivity. J Urol 181: 1459–1466, 2009.

19. Yew WP, Hibberd T, Spencer NJ, and Zagorodnyuk V. Piezo1, but not ATP, is required for mechanotransduction by bladder mucosal afferents in cystitis. Autonomic neuroscience : basic & clinical 256: 103231, 2024.

20. Ramsay S, Yew WP, Brookes S, and Zagorodnyuk V. A combination of peripherally restricted CB(1) and CB(2) cannabinoid receptor agonists reduces bladder afferent sensitisation in cystitis. Eur J Pharmacol 985: 177078, 2024.

21. Aizawa N, Fukuhara H, Fujimura T, Homma Y, and Igawa Y. Direct influence of systemic desensitization by resiniferatoxin on the activities of Aδ- and C-fibers in the rat primary bladder mechanosensitive afferent nerves. International journal of urology : official journal of the Japanese Urological Association 23: 952–956, 2016.

22. Daly D, Rong W, Chess-Williams R, Chapple C, and Grundy D. Bladder afferent sensitivity in wild-type and TRPV1 knockout mice. J Physiol 583: 663–674, 2007.

23. de Groat WC, and Yoshimura N. Afferent nerve regulation of bladder function in health and disease. Handbook of experimental pharmacology 91–138, 2009.

24. Yoshimura N, Erdman SL, Snider MW, and de Groat WC. Effects of spinal cord injury on neurofilament immunoreactivity and capsaicin sensitivity in rat dorsal root ganglion neurons innervating the urinary bladder. Neuroscience 83: 633–643, 1998.

25. Forrest SL, Osborne PB, and Keast JR. Characterization of bladder sensory neurons in the context of myelination, receptors for pain modulators, and acute responses to bladder inflammation. Frontiers in neuroscience 7: 206, 2013.

26. Sharma H, Kyloh M, Brookes SJH, Costa M, Spencer NJ, and Zagorodnyuk VP. Morphological and neurochemical characterisation of anterogradely labelled spinal sensory and autonomic nerve endings in the mouse bladder. Autonomic neuroscience : basic & clinical 227: 102697, 2020.

27. Hwang SJ, Oh JM, and Valtschanoff JG. The majority of bladder sensory afferents to the rat lumbosacral spinal cord are both IB4- and CGRP-positive. Brain Res 1062: 86–91, 2005.

28. Spencer NJ, Greenheigh S, Kyloh M, Hibberd TJ, Sharma H, Grundy L, Brierley SM, Harrington AM, Beckett EA, Brookes SJ, and Zagorodnyuk VP. Identifying unique subtypes of spinal afferent nerve endings within the urinary bladder of mice. J Comp Neurol 526: 707–720, 2018.

29. Usoskin D, Furlan A, Islam S, Abdo H, Lonnerberg P, Lou D, Hjerling-Leffler J, Haeggstrom J, Kharchenko O, Kharchenko PV, Linnarsson S, and Ernfors P. Unbiased classification of sensory neuron types by large-scale single-cell RNA sequencing. Nat Neurosci 18: 145–153, 2015.

30. Jung M, Dourado M, Maksymetz J, Jacobson A, Laufer BI, Baca M, Foreman O, Hackos DH, Riol-Blanco L, and Kaminker JS. Cross-species transcriptomic atlas of dorsal root ganglia reveals species-specific programs for sensory function. Nat Commun 14: 366, 2023.

31. Qi L, Iskols M, Shi D, Reddy P, Walker C, Lezgiyeva K, Voisin T, Pawlak M, Kuchroo VK, Chiu IM, Ginty DD, and Sharma N. A mouse DRG genetic toolkit reveals morphological and physiological diversity of somatosensory neuron subtypes. Cell 187: 1508–1526.e1516, 2024.

32. Sharma N, Flaherty K, Lezgiyeva K, Wagner DE, Klein AM, and Ginty DD. The emergence of transcriptional identity in somatosensory neurons. Nature 577: 392–398, 2020.

33. Fowler CJ, Griffiths D, and de Groat WC. The neural control of micturition. Nature reviews Neuroscience 9: 453–466, 2008.

34. Cheng CL, Liu JC, Chang SY, Ma CP, and de Groat WC. Effect of capsaicin on the micturition reflex in normal and chronic spinal cord-injured cats. Am J Physiol 277: R786–794, 1999.

35. Cheng CL, Ma CP, and de Groat WC. Effects of capsaicin on micturition and associated reflexes in rats. Am J Physiol 265: R132–138, 1993.

36. Maggi CA, and Conte B. Effect of urethane anesthesia on the micturition reflex in capsaicin-treated rats. J Auton Nerv Syst 30: 247–251, 1990.

37. Kuga N, Tanioka A, Hagihara K, and Kawai T. Modulation of afferent nerve activity by prostaglandin E2 upon urinary bladder distension in rats. Exp Physiol 101: 577–587, 2016.

38. Herrera GM, Pozo MJ, Zvara P, Petkov GV, Bond CT, Adelman JP, and Nelson MT. Urinary bladder instability induced by selective suppression of the murine small conductance calcium-activated potassium (SK3) channel. J Physiol 551: 893–903, 2003.

39. Montalbetti N, Rooney JG, Marciszyn AL, and Carattino MD. ASIC3 fine-tunes bladder sensory signaling. Am J Physiol Renal Physiol 315: F870–f879, 2018.

40. Kullmann FA, Daugherty SL, de Groat WC, and Birder LA. Bladder smooth muscle strip contractility as a method to evaluate lower urinary tract pharmacology. J Vis Exp e51807, 2014.

41. Dalghi MG, Ruiz WG, Clayton DR, Montalbetti N, Daugherty SL, Beckel JM, Carattino MD, and Apodaca G. Functional roles for PIEZO1 and PIEZO2 in urothelial mechanotransduction and lower urinary tract interoception. JCI Insight 6: 2021.

42. Ness TJ, and Gebhart GF. Colorectal distension as a noxious visceral stimulus: physiologic and pharmacologic characterization of pseudaffective reflexes in the rat. Brain Res 450: 153–169, 1988.

43. Christianson JA, and Gebhart GF. Assessment of colon sensitivity by luminal distension in mice. Nature protocols 2: 2624–2631, 2007.

44. Heppner TJ, Hennig GW, Nelson MT, and Herrera GM. Afferent nerve activity in a mouse model increases with faster bladder filling rates in vitro, but voiding behavior remains unaltered in vivo. Am J Physiol Regul Integr Comp Physiol 323: R682–r693, 2022.

45. Dalghi MG, Clayton DR, Ruiz WG, Al-Bataineh MM, Satlin LM, Kleyman TR, Ricke WA, Carattino MD, and Apodaca G. Expression and distribution of PIEZO1 in the mouse urinary tract. Am J Physiol Renal Physiol 317: F303–f321, 2019.

46. Marshall KL, Saade D, Ghitani N, Coombs AM, Szczot M, Keller J, Ogata T, Daou I, Stowers LT, Bönnemann CG, Chesler AT, and Patapoutian A. PIEZO2 in sensory neurons and urothelial cells coordinates urination. Nature 588: 290–295, 2020.

47. Carrisoza-Gaytan R, Mutchler SM, Carattino F, Soong J, Dalghi MG, Wu P, Wang W, Apodaca G, Satlin LM, and Kleyman TR. PIEZO1 is a distal nephron mechanosensor and is required for flow-induced K+ secretion. J Clin Invest 134: 2024.

48. Yang X, Zeng H, Wang L, Luo S, and Zhou Y. Activation of Piezo1 downregulates renin in juxtaglomerular cells and contributes to blood pressure homeostasis. Cell Biosci 12: 197, 2022.

49. Mikesell AR, Isaeva O, Moehring F, Sadler KE, Menzel AD, and Stucky CL. Keratinocyte PIEZO1 modulates cutaneous mechanosensation. eLife 11: 2022.

50. Cahalan SM, Lukacs V, Ranade SS, Chien S, Bandell M, and Patapoutian A. Piezo1 links mechanical forces to red blood cell volume. eLife 4: 2015.

51. Choi D, Park E, Jung E, Cha B, Lee S, Yu J, Kim PM, Lee S, Hong YJ, Koh CJ, Cho CW, Wu Y, Li Jeon N, Wong AK, Shin L, Kumar SR, Bermejo-Moreno I, Srinivasan RS, Cho IT, and Hong YK. Piezo1 incorporates mechanical force signals into the genetic program that governs lymphatic valve development and maintenance. JCI Insight 4: 2019.

52. Zhu T, Guo J, Wu Y, Lei T, Zhu J, Chen H, Kala S, Wong KF, Cheung CP, Huang X, Zhao X, Yang M, and Sun L. The mechanosensitive ion channel Piezo1 modulates the migration and immune response of microglia. iScience 26: 105993, 2023.

53. Hill RZ, Loud MC, Dubin AE, Peet B, and Patapoutian A. PIEZO1 transduces mechanical itch in mice. Nature 607: 104–110, 2022.

54. Heppner TJ, Tykocki NR, Hill-Eubanks D, and Nelson MT. Transient contractions of urinary bladder smooth muscle are drivers of afferent nerve activity during filling. J Gen Physiol 147: 323–335, 2016.

55. Thorneloe KS, Knorn AM, Doetsch PE, Lashinger ES, Liu AX, Bond CT, Adelman JP, and Nelson MT. Small-conductance, Ca(2+) -activated K+ channel 2 is the key functional component of SK channels in mouse urinary bladder. Am J Physiol Regul Integr Comp Physiol 294: R1737–1743, 2008.

56. Herrera GM, Etherton B, Nausch B, and Nelson MT. Negative feedback regulation of nerve-mediated contractions by KCa channels in mouse urinary bladder smooth muscle. Am J Physiol Regul Integr Comp Physiol 289: R402–r409, 2005.

57. Mills KA, West EJ, Grundy L, McDermott C, Sellers DJ, Rose’Myer RB, and Chess-Williams R. Hypersensitivity of bladder low threshold, wide dynamic range, afferent fibres following treatment with the chemotherapeutic drugs cyclophosphamide and ifosfamide. Arch Toxicol 94: 2785–2797, 2020.

58. Delmas P, Hao J, and Rodat-Despoix L. Molecular mechanisms of mechanotransduction in mammalian sensory neurons. Nature reviews Neuroscience 12: 139–153, 2011.

59. Saal HP, Delhaye BP, Rayhaun BC, and Bensmaia SJ. Simulating tactile signals from the whole hand with millisecond precision. Proc Natl Acad Sci U S A 114: E5693–e5702, 2017.

60. Vlaskovska M, Kasakov L, Rong W, Bodin P, Bardini M, Cockayne DA, Ford AP, and Burnstock G. P2X3 knock-out mice reveal a major sensory role for urothelially released ATP. J Neurosci 21: 5670–5677, 2001.

61. Cockayne DA, Dunn PM, Zhong Y, Rong W, Hamilton SG, Knight GE, Ruan HZ, Ma B, Yip P, Nunn P, McMahon SB, Burnstock G, and Ford AP. P2X2 knockout mice and P2X2/P2X3 double knockout mice reveal a role for the P2X2 receptor subunit in mediating multiple sensory effects of ATP. J Physiol 567: 621–639, 2005.

62. Cockayne DA, Hamilton SG, Zhu QM, Dunn PM, Zhong Y, Novakovic S, Malmberg AB, Cain G, Berson A, Kassotakis L, Hedley L, Lachnit WG, Burnstock G, McMahon SB, and Ford AP. Urinary bladder hyporeflexia and reduced pain-related behaviour in P2X3-deficient mice. Nature 407: 1011–1015, 2000.

63. Rong W, Spyer KM, and Burnstock G. Activation and sensitisation of low and high threshold afferent fibres mediated by P2X receptors in the mouse urinary bladder. J Physiol 541: 591–600, 2002.

64. Takezawa K, Kondo M, Kiuchi H, Ueda N, Soda T, Fukuhara S, Takao T, Miyagawa Y, Tsujimura A, Matsumoto-Miyai K, Ishida Y, Negoro H, Ogawa O, Nonomura N, and Shimada S. Authentic role of ATP signaling in micturition reflex. Sci Rep 6: 19585, 2016.

65. Birder L, and Andersson KE. Urothelial signaling. Physiol Rev 93: 653–680, 2013.

